# Intraspecific variation in migration timing of green sturgeon in the Sacramento River system

**DOI:** 10.1101/2021.12.03.471146

**Authors:** Scott F. Colborne, Lawrence W. Sheppard, Daniel R. O’Donnell, Daniel C. Reuman, Jonathan A. Walter, Gabriel P. Singer, John T. Kelly, Michael J. Thomas, Andrew L. Rypel

**Author notes:** **Corresponding Author:** Scott F. Colborne, Department of Wildlife, Fish, and Conservation Biology, University of California, Davis, CA USA. Denotes co-lead authors (Scott F. Colborne, Lawrence W. Sheppard).

## Abstract

**Background:** Understanding movement patterns of anadromous fishes is critical to conservation management of declining wild populations and preservation of habitats. Yet, infrequent observations of individual animals fundamentally constrain accurate descriptions of movement dynamics.

**Methods:** In this study, we synthesized over a decade (2006–2018) of acoustic telemetry tracking observations of green sturgeon (*Acipenser medirostris*) in the Sacramento River system to describe major anadromous movement patterns.

**Results:** We observed that green sturgeon exhibited a unimodal in-migration during the spring months but had a bimodal distribution of out-migration timing, split between an ‘early’ out-migration (32%) group during May - June, or alternatively, holding in the river until a ‘late’ out-migration (68%), November - January. Focusing on these out-migration groups, we found that river discharge, but not water temperature, may cue the timing of migration, and that fish showed a tendency to maintain out-migration timing between subsequent spawning migration events.

**Conclusions:** We recommend that life history descriptions of green sturgeon in this region reflect the distinct out-migration periods described here. Furthermore, we encourage the continued use of biotelemetry to describe migration timing and life history variation, not only this population but other green sturgeon populations and other species.

## Background

Humans modify waterways to suit energy, industrial, agricultural, and drinking water needs, and these modifications impact the life-histories of migratory fishes. For example, dams and diversions change river flow patterns and often reduce discharge rates in natural channels, present direct barriers to fish movements, and cause loss of essential feeding, spawning, and nursery habitats (1–3). Yet poor understanding of species life-histories often results in infrastructure that fundamentally blocks migrations or are otherwise not wildlife-friendly (4, 5). As a result, conservation and management efforts often focus on restoring functionality to riverine systems by removing barriers or through modifications that provide for fish passage around barriers (6, 7). These efforts are attempts to balance human needs (i.e., ecosystem services) with the ecological requirements of the other organisms that depend on these systems (7).

Life history strategies vary not only among but also within species, with both inter- and intraspecific diversity contributing to community structure and stability (8, 9). Intraspecific variation may buffer populations from stochastic events that affect certain locations or time periods, e.g., through portfolio effects (10–12). Chinook salmon (*Oncorhynchus tshawytscha*) can broadly be described as migrating to oceans as smolts and spending several years growing and maturing in salt water before returning to their natal streams and rivers (13). However, Chinook salmon also exhibit variation in migration timing of adults and smolts (14–16). In comparison to the temporal variability in Chinook salmon migration, striped bass (*Morone saxatilis*) in the Hudson River Estuary migrate different distances to spawning sites and maintain these site preferences across years (17), effectively dispersing reproductive effort across multiple sections of the river. Both temporal and spatial intraspecific variation have implications for the populations involved and fisheries that use this natural resource. Effective management of wild populations requires understanding diversity in migratory tactics, but intraspecific variation may be hard to document while monitoring wild fishes.

Along the Pacific coast of North America, green sturgeon (*Acipenser medirostris*) are a long-lived, intermittent spawning fish of conservation concern. The southern distinct population segment (sDPS) in the Sacramento River of California is listed as ‘threatened’ under the US Endangered Species Act (ESA), and throughout their range, green sturgeon are listed under CITES (Convention on International Trade in Endangered Species of Wild Fauna and Flora) Appendix II (https://cites.org) (18, 19). A conceptual model of sDPS green sturgeon movements described adult green sturgeon migrations as following these general steps: (1) in-migration to the river occurs during the spring months, peaking in March, with fish travelling over 400 km upriver to spawning grounds; (2) spawning occurs during the late spring months (April through June); (3) adults then spend the summer months in the river near to spawning grounds; (4) out-migration to the Pacific Ocean happens over an extended period in the late summer through autumn months; (5) individuals then remain in the Pacific Ocean for 2–4 years between migration events (see 20 for details). Conceptual models like these are valuable for general life history descriptions of wild animals but can be improved upon when high-resolution data on movements of individuals become available.

The specific objectives of this study were to synthesize the long-term movement profiles of individual green sturgeon from acoustic telemetry to (1) describe timing of green sturgeon migrations in the Sacramento River; (2) determine if swim-down events were correlated with environmental variables (discharge and temperature); and (3) evaluate whether individuals exhibited fidelity to a particular pattern in migration timing across migration events. We expected upriver migrations in the spring months would occur over a single period so that adult green sturgeon arrived at spawning grounds when conditions were optimal for the development of eggs and larvae. We did not have similar expectations for out-migration movements. Past observations of green sturgeon have described out-migrations occurring across a span of several months (21, 22), including two pulses of downriver movements in the Klamath and Trinity rivers (northern distinct population segment, see 23).

## Materials & Methods

### Study system and acoustic tagging

The lower Sacramento River system of California, defined here as the portion of the river below the Keswick dam (40.612°, -122.446°), is over 400 km long and is fed by multiple tributaries before it joins the San Francisco Estuary and the Pacific Ocean. This is among the planet’s most human-altered river systems: flow pathways, discharge rates, and fish habitat are all subject to human control (24), and the Sacramento River system remains an area of great conservation interest. The Sacramento River system is perhaps most widely known for its Chinook salmon (*Oncorhynchus tshawytscha*; e.g., 25) and steelhead trout (*Oncorhynchus mykiss*; e.g., 26), but it is also the central hub for a rich and highly endemic assemblage of fishes in California (27). It is also a highly threatened fauna, with an estimated 83% of species currently classified as in some form of decline (28).

Over 300 acoustic receivers (VR2W-69 kHz, Innovasea Inc., Halifax NS, Canada; Fig. 1) have been deployed throughout the Sacramento River, the Inland Delta, the San Francisco Estuary, and nearshore regions of the Pacific Ocean to passively monitor the movements of tagged fish (see also 21,22). During the lifetime of these acoustic telemetry projects, deployment configurations and receiver coverage have varied to address the questions of specific projects. However, there was consistent coverage throughout much of the Sacramento River system during the 12-year observation period (2006-2018) used in this study.

**Figure 1.**
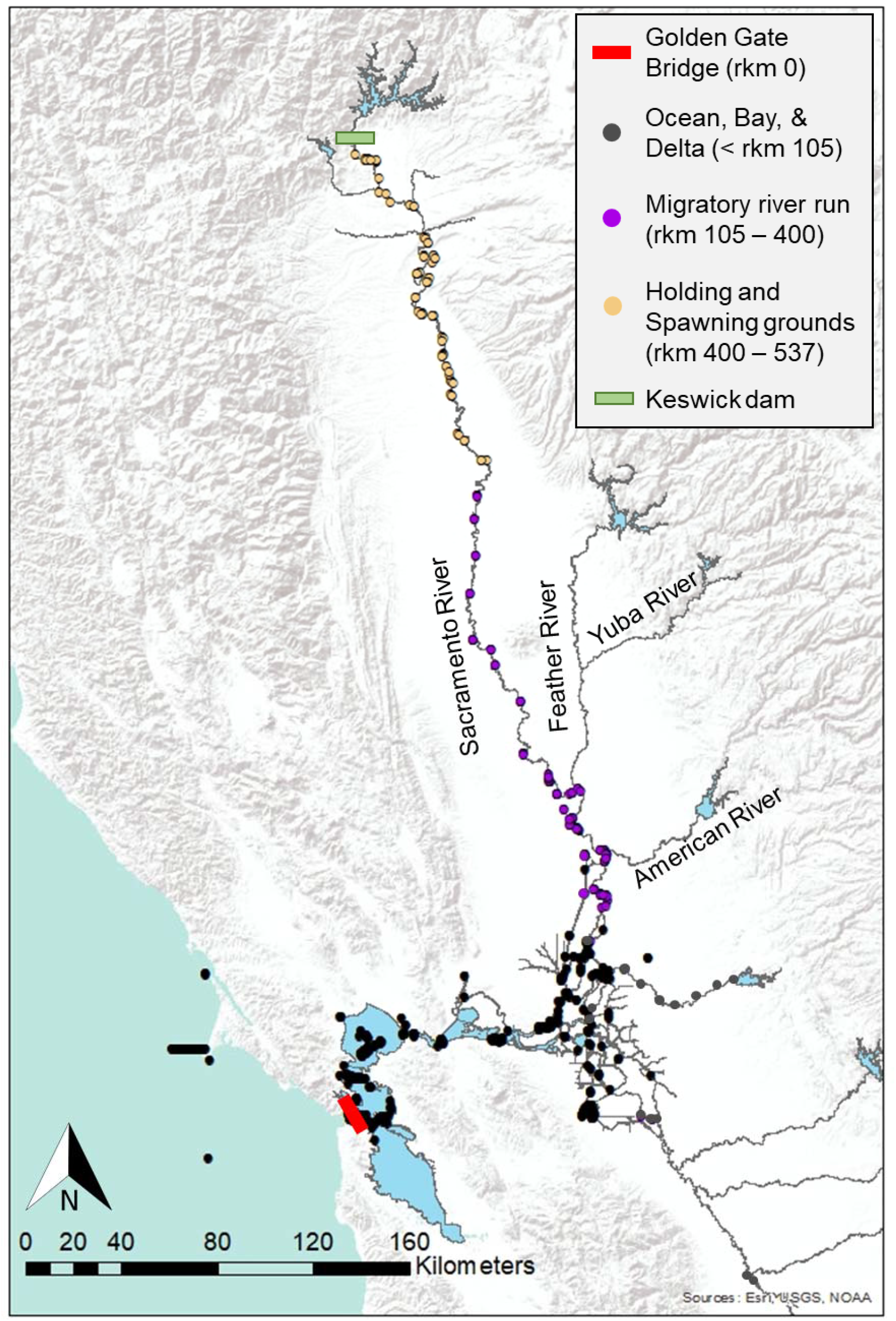
Positions of receivers throughout the San Francisco Bay, Sacramento River, and San Joaquin River systems during the 2006 through 2018 observation period used in this study.

From 2002 to 2014, green sturgeon were surgically implanted with 10-year lifespan acoustic transmitters (n = 350 tagged sturgeon considered for analysis) by scientists from 13 agencies and research groups (21,22,29,30). Tagging and capture methods varied slightly across agencies and research groups and over time but were consistent in the following respects. Fish tagged and released were caught by trammel or gillnets at multiple locations in the Sacramento River, San Pablo Bay, and Suisun Bay. Nets were either watched for movement in float lines or soaked for a maximum of 30 min to minimize capture stress on green sturgeon. Fish selected for tagging were inverted to initiate a tonic immobility-like state and water flow was provided continuously across their gills. The surgical procedure for these fish is outlined in Miller et al. (22) but is reviewed briefly here. A small incision, approximately 30 mm, was made anterior to the pelvic girdle and offset from the midline. Green sturgeon were tagged with V16 (n = 322), V13 (n = 5), or V9 (n = 23) transmitters, depending on the size of the fish being tagged (Innovasea Inc., Halifax, NS Canada). Transmitters were inserted into the body cavity and the incision closed with two to three simple interrupted sutures (PDS II Violet Monofilament, absorbable) tied with 2×2 surgeon’s knots. This general surgical method has been employed successfully on multiple fishes, including sturgeon, and has been shown to not impact survival, growth, or swimming performance of juvenile green sturgeon (31). In total, the surgical period lasted approximately 5 minutes, after which the fish were immediately released at the site of capture or, if showing signs of stress, temporarily held in an aerated stock tank until they resumed normal responses.

### Data selection and longitudinal profiles

The Interagency Telemetry Advisory Group (ITAG) within the Interagency Ecological Program (IEP), formerly California Fish Tracking Consortium (CFTC), has coordinated the telemetry efforts of multiple agencies and researchers throughout the Central Valley of California. As part of the research coordination effort, the BARD database established through the UC Davis Biotelemetry Lab (http://cftc.metro.ucdavis.edu/biotelemetry-autonomous-real-time-database/landingmap) has been used to compile detections records across a greater scale than a single organization could compile. The BARD data repository has provided spatial and temporal resolution to facilitate synthesis-based studies of green sturgeon across multiple projects and tagging efforts. We accessed green sturgeon detection data from 2006 through 2018 from the BARD database (see above). For individual green sturgeon, we compiled detection data across the Sacramento River system to create an ordered series of detections that included time stamps of detection, location of receivers recording each detection, and river km (measured as distance from the entrance to the Pacific Ocean marked by the Golden Gate Bridge in San Francisco Bay, hereafter ‘rkm’).

Green sturgeon longitudinal detection profiles based on rkm were used to determine when fish were moving into and out of the river system (see Fig. 1 for general area categories). When a fish was first detected downriver and then upriver in the same calendar year, the date of the first record above rkm 105 was logged as the ‘up-date’ and considered when the fish began migration upriver, presumably to spawn. We selected the 105 rkm value to reflect transition from Suisun Marsh and the inland Delta region into the Sacramento River proper (Fig. 1). When a fish was detected upriver in a calendar year and then detected downriver within 500 days of the beginning of that calendar year, the date of first record below rkm 400 was classified as the ‘down-date’ and recorded as the day outmigration commenced. We selected a value of 400 rkm as the threshold separating spawning and holding grounds from the rest of the river based on visual inspection of movement paths (see Supplemental Materials Fig. S1) and previously used thresholds (32) based on descriptions of green sturgeon activity in the Sacramento River (20).

To describe the distribution of days fish either began moving up the Sacramento River or began their out-migration back down the river, a cumulative distribution function tallying the observed event dates (see above) was used. If we observed a plateau in dates consistent across observation years, i.e., a period during which no migration events are observed, the mid-value of the plateau was used as a dividing value between groups.

Because green sturgeon migrations often spanned more than one calendar year, day of the year alone could not be used to measure migration timing. Rather, we tabulated day counts relative to the year that a green sturgeon began upriver migration runs, e.g., a fish that migrated upriver on 1 March, 2017 and downriver on 5 January, 2018 would have been described as moving upriver on Day 60 and downriver on Day 370. Dates calculated using this method are referred to as ‘journey dates’ and were all calculated relative to 1 January of the year that a green sturgeon began an upriver migration.

Many green sturgeon were tagged in the Sacramento River during spring months, presumably already on a spawning run up the river at time-of-tagging, with 71 sturgeon tagged between rkm 150 and 518. For these fish, the first detected migration event using the criteria outlined above was their downriver migration. As such, the greater number of downriver migration events observed (n = 224 events) as compared to upriver migration events (n = 129 events) was attributed primarily to tagging in the river. We established upriver and downriver migration dates representing the complete history of a migration event, referred to as ‘paired up-down dates’ for 117 migration events during the observation period.

### Environmental data

We collated environmental data from multiple long-term monitoring stations in the Sacramento River. We compiled upriver environmental data spanning the study period from two stations: discharge rates were retrieved from records of the ORD station (California Department of Water Resources, National Wildlife Refuge Ord Bend Unit; 39.628°, -121.993°) and temperature data from the RDB station (US Forest Service, Red Bluff Recreation Area; 40.154°, -122.202°). These stations are located ∼60 km apart but chosen for having the most complete records covering the observation period and their proximity to upriver habitats used by green sturgeon. For downriver sites, we collected data from two stations: discharge rates from the DTO station (California Department of Water Resources, Delta out discharge station; 38.059°, - 122.025°) and temperature data from the RVB station (California Department of Water Resources, Rio Vista Bridge; 38.160°, -121.686°), which were approximately 30 km apart.

### Environmental variables and migrations

For each up- or down-river migration event that could be determined from acoustic detections, we created profiles of discharge and temperature for the 21 days surrounding the day of migration. Each date that a migration was deemed to have commenced (either upriver or downriver) was designated Day 0; we then extracted river discharge and temperature data from the 14 d prior to that date and the 7 d afterwards to form a profile around each migration event. A corresponding segment of environmental data (discharge and temperature) was extracted from the matched dates using the stations outlined above, i.e., DTO/RVB for upriver and ORD/RDB for downriver profiles. Once profiles were created for individual fish, the mean values across all fish for discharge and temperature in each of the 22-days was determined to provide a qualitative description of these parameters in relation to migration events.

We had *a priori* interest in downriver migrations due to the extended total duration over which they have been previously observed to occur. We therefore used semiparametric Cox proportional hazard regression (hereafter CPH) (33) to determine whether temperature and/or discharge characteristics were predictive of downriver migrations. Environmental parameters and swim down events were binned into 5-d intervals. The covariates of mean temperature and discharge were tested for collinearity (34) during the ‘early’ and ‘late’ out-migration periods based on variance inflation factor (VIF) scores. Both the ‘early’ and ‘late’ out-migration groups (VIF≤1.17) were found to be below the collinearity threshold for collinearity of 5.0 (34, 35). Within each 5-d interval, we determined the mean discharge (m^3^ s^-1^), minimum daily discharge (m^3^ s^-1^), and mean water temperature (°C). We also considered the percent change in mean discharge (Δ discharge) and percent change in mean water temperature (Δ temperature) calculated as the change between subsequent time intervals, i.e., ((Temp_(t+1)_ – Temp_(t)_) / Temp_(t)_) x 100 and ((Discharge_(t+1)_ – Discharge_(t)_) / Discharge_(t)_) x 100.

The CPH model can be described by the formula

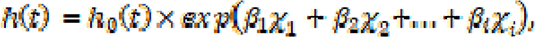

where *h(t)* is the “expected hazard” at time t, *h_0_(t)* represents the “baseline hazard” assuming no effect of any covariates, and β*_i_* is the regression coefficient for an explanatory variable (χ*_i_*). A hazard ratio (HR) independent of time (*t*) was estimated for each explanatory variable (χ*_i_*) based on the regression coefficients determined in the CPH model 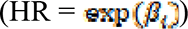. A positive regression coefficient, and consequently HR value > 1, for a covariate indicates a positive relationship between that covariate and, in this case, greater migration probability; larger values of HR indicate a greater probability of migration for each unit increase in the explanatory variable. The CPH models were fitted using the *coxph* function in the *survival* analysis R package (36). The proportional hazard for each covariate is assumed, by the CPH approach, to be independent of time; this was tested using the Schoenfeld residuals (*cox.zph* function), with p > 0.05 for each covariate in the model considered passing this assumption. Due to violations of this assumption when including migration year as a continuous variable, we included year as a stratification factor.

We used an information-theoretic model selection process that considered models including all possible combinations of the five covariates outlined above. Using Akaike’s Information Criterion (AIC), the model with the lowest AIC score was identified as the most supported model (ΔAIC = 0), but all models with Δ AIC < 4 were considered competitive (Arnold 2010). The relative importance for each covariate was determined by summing the AIC weights of the models in which they occurred. CPH model coefficients were determined by using model averaging across the 90% confidence set of models (37, 38) using the *dredge* and *model.avg* functions of *MuMIn* R package (39).

### Repeatability in timing of migration events

Due to the duration of monitoring covered in this study, some fish made more than one migration up and down the river. Where multiple migration events occurred for the same fish, it was possible to compare the first and second swim down dates to determine if the timing of migration was correlated across events. To determine if there was repeatability in the timing of migrations, journey days were correlated between (i) first and second upriver migrations, (ii) first and second downriver migrations, and (iii) upriver and downriver migrations. If individuals were observed making more than two migrations during the observation period, only the first and second event were considered. Normality was tested with the Shapiro-Wilk test. If normality was violated (P < 0.05) for either of the two variables for which correlation was to be computed, Spearman’s rank correlation was used, otherwise Pearson’s correlation was used. For correlation testing purposes, only paired migration events with both up and down dates observed were considered because up and down movement dates were involved in the comparisons; this constraint was applied for the up/up and down/down comparisons, as well as for the up/down comparisons, to produce a consistent dataset on which all the correlation results were based.

Given the presence of two distinct out-migration groups (see Results), we evaluated if the number of fish observed switching between groups from their first to their second trips down the river was more compatible with consistency or randomness through time of individual fish departure timing. A randomization trial performed in MatLab (version R2021a, updated 13 April 2021) was used to test the null hypothesis that fish selected their downriver migration group at random and that the out-migration timing during the second event was not related to the first event. For each simulation, the same number of fish observed in the field making multiple migration events (n = 64) were randomly assigned to either an ‘early’ (journey day < 250) or ‘late’ (journey day > 250) out-migration group (first migration), and then independently assigned to a second ‘early’ or ‘late’ group (second migration). Probabilities for assignments were based on empirical values. The number of fish that switched out-migration timing, i.e., ‘early-to-late’ or ‘late-to-early’, was tallied to determine the total number of switches that occurred in the simulation. This process was repeated for 100,000 simulations to generate a distribution of numbers of migration timing switches that would be expected if migration timing was random and independent across events. The empirical number of switches was then compared to the distribution of simulated values, and if the observed number was within the 95% confidence interval of the simulated values, fish were considered to show significant consistency, through time, in their migration group choice. All fish were included in this analysis for which at least two downriver dates were available.

To determine if river discharge was related to repeatability in the migration timing of fish, we first determined if there was a linear relationship between the journey-day departure timing of first and second out-migration events for fish that departed ‘early’ for both migrations (i.e., ‘early-early’ fish). Because there was evidence that ‘early-early’ fish out-migration journey days were strongly correlated between migrations (see Results), the out-migration dates for fish that switched groups between events (i.e., ‘early-to-late’, n = 13, and ‘late-to-early’, n = 8) were used to estimate matched dates for the ‘early’ period for the year they migrated ‘late’ based on the regression relationship determined for ‘early-early’ fish. For each of the fish that switched migration groups this provided an observed ‘early’ migration date (either first or second migration) and regression-inferred ‘early’ date (i.e., for the year the individual was observed migrating ‘late’). Mean flow over a 7-day period leading up to the out-migration dates was determined for the years fish were observed departing ‘early’ and compared to the years in which they instead held over and departed as part of the ‘late’ group. Discharge was compared between the two groups (observed ‘early’ departure and regression-matched ‘early’ dates) using a paired Wilcoxon rank sums test. Here, we systematically evaluated river flow during actual ‘early’ departures compared to matched ‘early’ times when a fish could, in principle, have departed but instead chose to migrate ‘late’.

## Results

Based on the tagging records of 350 individual green sturgeon, 151 individuals were detected in the Sacramento River system during 2006–2018. Nine fish were tagged prior to the beginning of the observation period (2003–2005), while the remainder of green sturgeon were tagged 2006–2013 (2006, n = 21; 2008, n = 10; 2009, n = 29; 2010, n = 18; 2011, n = 46; 2012, n = 16; 2013, n = 2). Average fork length of green sturgeon tagged (mean ± 1 S.E.) was 1726 ± 16 mm. Fish of both sexes were represented in the sample (females = 15, males = 35), but most green sturgeon were not sexed (n = 101) during tagging due to logistical and permitting constraints.

A total of 129 upriver migrations and 224 downriver movements were identified (Table 1), including 117 paired detection events (n = 85 unique sturgeon) covering the full migration sequence (i.e., migration upriver to spawning grounds and subsequent out-migration back downriver to the Pacific Ocean). When considering all downriver movements including migration events without a corresponding upriver date, there were 62 green sturgeon that made more than one downriver migration during the observation period with a mean interval between downriver migration events of 4.3 years (1562 days).

**Table 1.** Counts of the numbers of unique green sturgeon observed undertaking upriver and downriver migrations in each calendar year (2006 – 2018). See methods for criteria used to identify that individual green sturgeon were migrating through the Sacramento River system in each year.

| **Year** | **Count of upriver**<br>**migrating fish** | **Count of downriver**<br>**migrating fish** |
| --- | --- | --- |
| 2006 | 2 | 18 |
| 2007 | 4 | 3 |
| 2008 | 2 | 8 |
| 2009 | 3 | 20 |
| 2010 | 4 | 34 |
| 2011 | 3 | 13 |
| 2012 | 18 | 36 |
| 2013 | 14 | 5 |
| 2014 | 14 | 24 |
| 2015 | 20 | 19 |
| 2016 | 28 | 28 |
| 2017 | 17 | 10 |
| 2018 | 0 | 6 |
| **Total** | **129** | **224** |

### Timing and duration of migration

Across all observation years, the 117 paired migration events began with green sturgeon swimming into the Sacramento River on 22 Mar ± 22 days (range 10 Feb – 14 Jun) (see Table 2 for individual year summaries). Across all fish, mean date of outmigration was 16 Oct ± 93 days (min. 15 Apr, max. 24 Mar of year following upriver migration) and individual green sturgeon were present in the Sacramento River for an average of 204 ± 97 days (Fig. 2a). Upriver migration dates followed a unimodal distribution that peaked in late Mar to early Apr (Fig. 2b). In comparison, downriver migration dates followed a bimodal distribution (Fig. 2c), with some green sturgeon returning downriver during May – Jun, and others remaining for several months, out-migrating to the Pacific Ocean during Nov – Jan (Table 2). All tagged green sturgeon returned to the Pacific Ocean after a holding period, and there was no evidence of permanent river-residency by any tagged fish.

**Figure 2.**
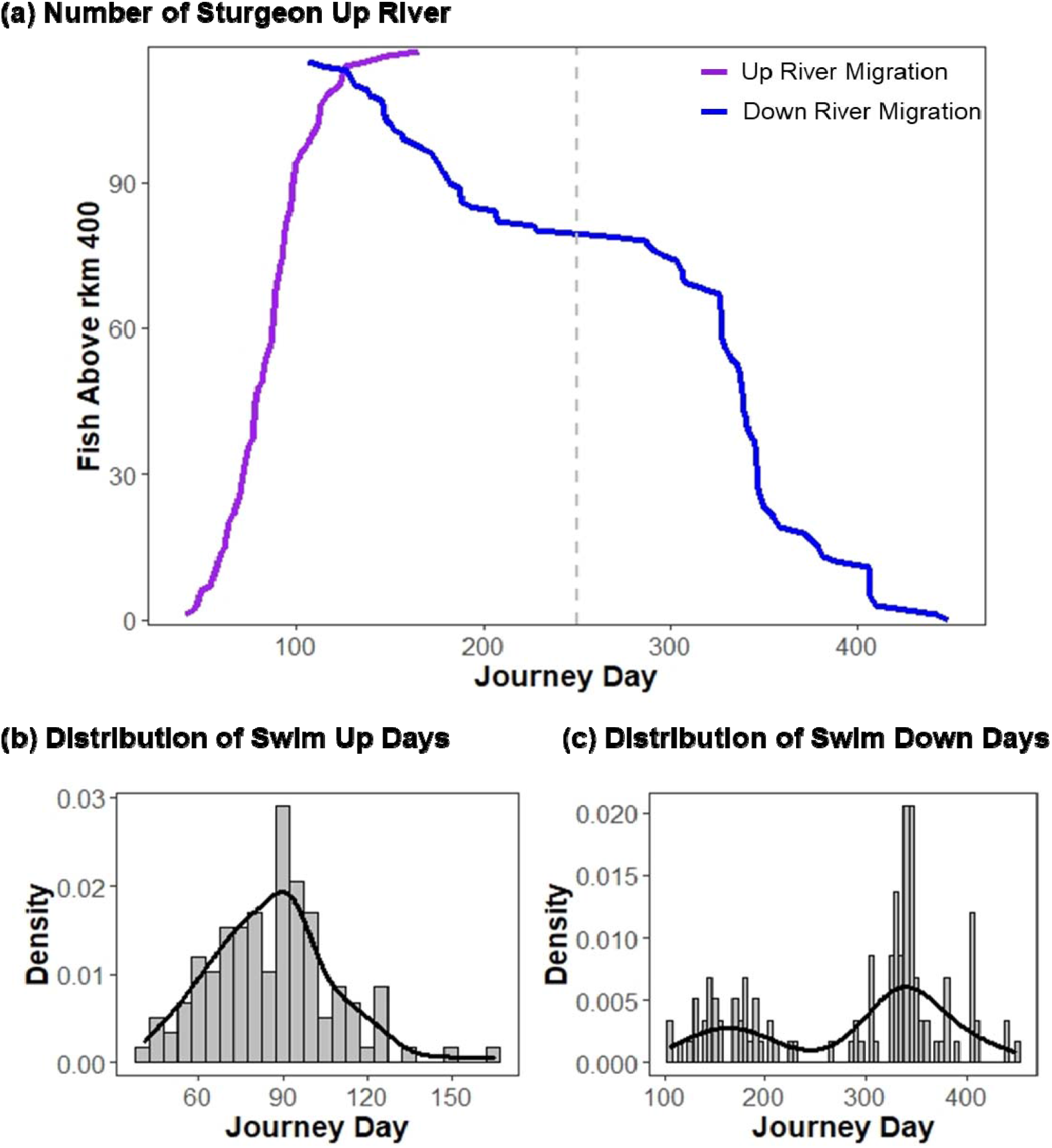
Cumulative number of fish migration events (max = 117) commenced by a given day of calendar year (journey day). Upriver migration (red) were used to determine the calendar year migrations began and this start year was applied to downriver migrations (blue), therefore, journey days > 365 indicate a fish that migrated upriver in one calendar year, e.g., 2013, and migrated downriver the following calendar year, e.g., 2014. The distribution of (b) swim up and (c) swim down dates are shown relative to a normal distribution.

**Table 2.**
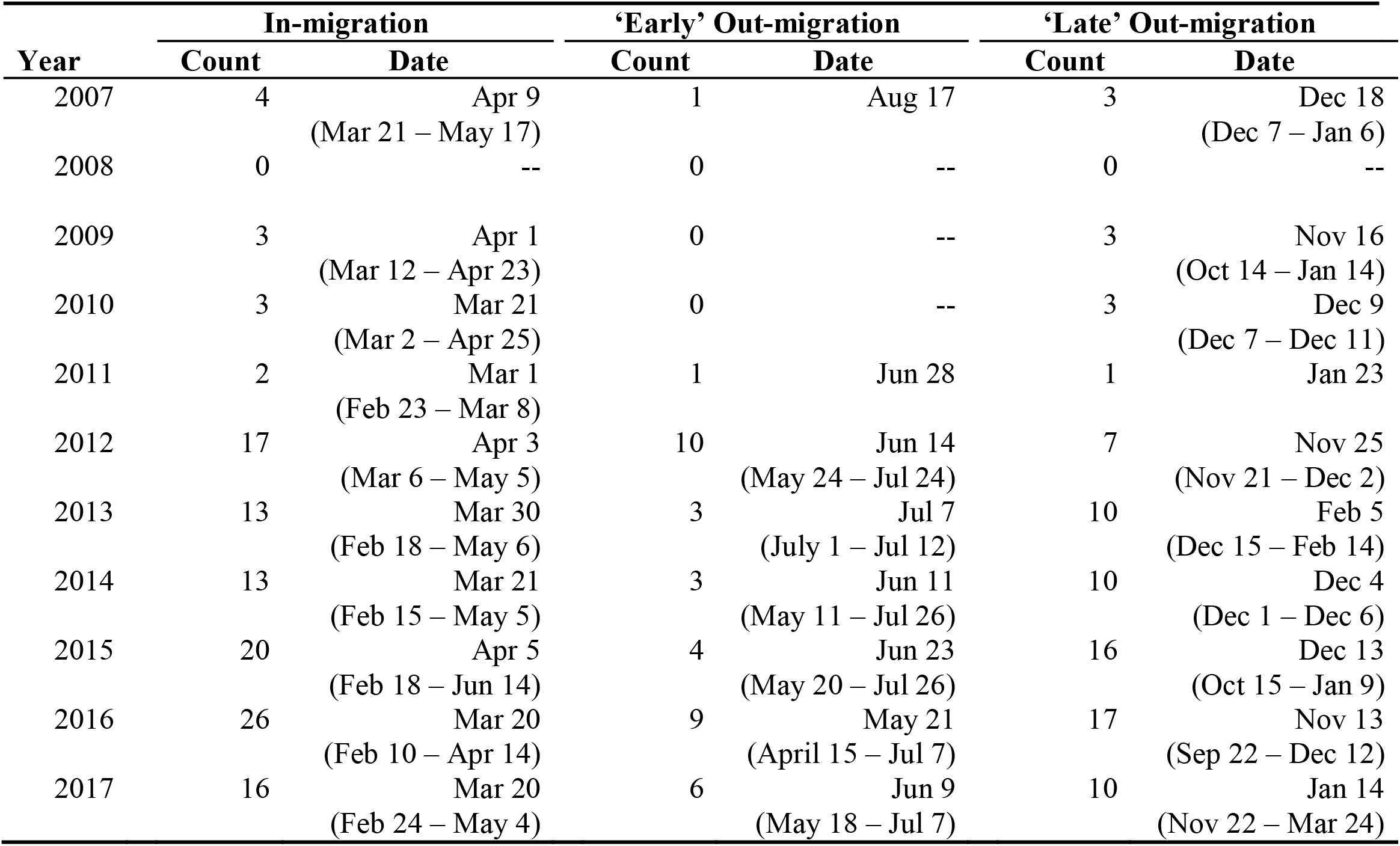
Count and dates of migrations for green sturgeon divided into in-migration, ‘early’ out-migration, and ‘late’ out-migration groups. The mean and range (in parentheses) of dates are presented based on the year a fish was detected migrating up the Sacramento River.

Based on the distributions of downriver migration dates we defined two swim-down groups, ‘early’ fish migrated downriver before Sep (< day 250) and ‘late’ fish spent a period of several extra months in the upper Sacramento River before returning to the Pacific Ocean during late autumn and early winter (> day 250; Fig. 2). Across the 117 paired up-down migration events, we observed 37 early return events (n = 34 unique individuals) that spent 76 days in the river system from swim up to swim down (range 26 – 142 days). In comparison, there were 80 late return events (n = 71 unique individuals) with fish spending a mean of 263 days in the river (range 139 – 368 days).

‘Early’ returning green sturgeon began their downriver migration on 12 Jun ± 31 days (min. 15 Apr, max. 17 Aug) and ‘late’ returning fish began their migration on 14 Dec ± 37 days (min. 22 Sep, max. 24 Mar of the year following migration). The days ‘early’ out-migration began occurred at higher water temperatures and lower discharge levels (mean ± 1 S.E., interquartile range; temperature: 14.88 ± 0.22 °C, IQR = 2.11 °C; discharge: 227.66 ± 11.20 m^3^ s^-1^, IQR = 119.28 m^3^ s^-1^) as compared to the ‘late’ out-migration group (temperature: 11.20 ± 0.18 °C, IQR = 2.32 °C; discharge: 346.00 ± 21.32 m^3^ s^-1^, IQR = 195.12 m^3^ s^-1^) (see Fig. S2 and S3 for further details).

### Environmental variables and migrations

Based on the unimodal distribution of upriver migrations and bimodal distribution of downriver dates described above, we described green sturgeon as migrating ‘upriver’ in a single group and out-migrating in two distinct groups based on timing (‘early’ and ‘late’). We created 22-day profiles relating discharge and water temperature to each of these three movement groups. Based on these profiles, green sturgeon began upriver movements during periods of increasing temperature as winter concluded (Fig. 3a). There was no visually detectable pattern in discharge and timing of downriver migration in the ‘early’ out-migration group (Fig. 3b), but there was a tendency for the ‘late’ out-migration group to begin downriver movements after an initial increase in discharge (Fig. 3c). Downriver migrations did not have apparent correlations to water temperature.

**Figure 3.**
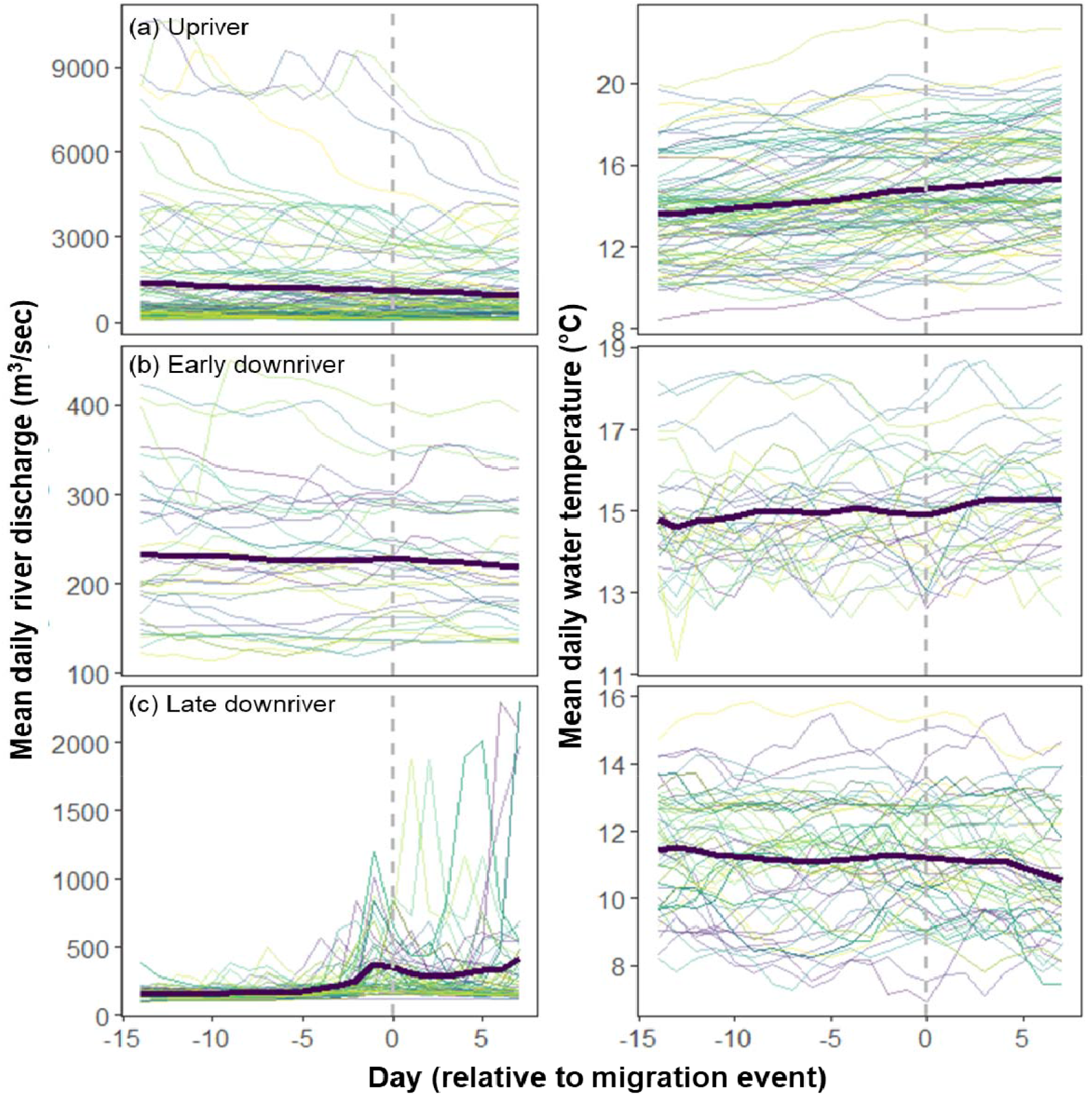
Profiles of Sacramento River discharge rate (m^3^ s^-1^) and temperature (°C) over a 21-day period surrounding the day of migration (14 days before and 7 days following migration). Profiles were created for (a) upriver migration dates, (b) ‘early’ downriver migrations, and (c) ‘late’ downriver migrations. Day 0 (dashed line) represents the date identified as the beginning of out-migration. The black line in each panel represents the mean discharge rate or temperature across all fish for each day and each colored line tracks an individual fish over 22 days. Environmental measures were collected from two stations each for the upriver and downriver migrations.

There were ten competitive models for the ‘early’ out-migration period (ΔAIC≤2.78) and 15 for the ‘late’ out-migration period (ΔAIC ≤3.90; Table 3). Relative importance among possible covariates in the ‘early’ group was highest for minimum discharge (0.98), followed by Δ discharge (0.73), mean discharge (0.46), Δ temperature (0.36), and was lowest for mean temperature (0.34). In the ‘late’ out-migration group, relative importance was highest for minimum discharge (0.93), followed by Δ discharge (0.55), mean discharge (0.44), mean temperature (0.43), and Δtemperature (0.42).

**Table 3.** Model fit summary for combinations of five covariations related to water discharge and temperature predicted to be related to the migration timing of (a) ‘early’ and (b) ‘late’ out-migration green sturgeon from the Sacramento River system. Predictor variables for each model are shown along with the number of parameters in each model (*K*), log likelihood (*LL*), Akaike’s Information Criterion (AIC), difference in AIC score compared to the top model (Δ*AIC*), and model weight (*w_i_*). Models up to the first with a Δ*AIC* score > 4 are shown.

| Model variables | $K$ | $LL$ | $AICc$ | $\Delta AIC$ | $w_i$ |
| --- | --- | --- | --- | --- | --- |
| **(a) ‘Early’ out-migration** |  |  |  |  |  |
| $\Delta$ discharge + min discharge | 2 | -29.52 | 63.06 | 0 | 0.19 |
| $\Delta$ discharge + min discharge + $\Delta$ temp | 3 | -28.93 | 63.92 | 0.86 | 0.12 |
| $\Delta$ discharge + min discharge + temp | 3 | -29.00 | 64.07 | 1.01 | 0.11 |
| discharge + $\Delta$ discharge + min discharge | 3 | -29.14 | 64.35 | 1.28 | 0.10 |
| discharge + min discharge | 2 | -30.39 | 64.80 | 1.74 | 0.08 |
| discharge + $\Delta$ discharge + min discharge + $\Delta$ temp | 4 | -28.62 | 65.34 | 2.28 | 0.06 |
| discharge + $\Delta$ discharge + min discharge + temp | 4 | -28.65 | 65.40 | 2.34 | 0.06 |
| discharge + min discharge + $\Delta$ temp | 3 | -29.70 | 65.46 | 2.40 | 0.06 |
| discharge + min discharge + temp | 3 | -29.82 | 65.69 | 2.63 | 0.05 |
| $\Delta$ discharge + min discharge + temp + $\Delta$ temp | 4 | -28.87 | 65.84 | 2.78 | 0.05 |
| discharge + min discharge + temp + $\Delta$ temp | 4 | -29.54 | 67.17 | 4.10 | 0.02 |
| **(b) ‘Late’ out-migration** |  |  |  |  |  |
| $\Delta$ discharge + min discharge | 2 | -105.51 | 215.04 | 0 | 0.13 |
| $\Delta$ discharge + min discharge + temp | 3 | -104.80 | 215.62 | 0.58 | 0.10 |
| $\Delta$ discharge + min discharge + $\Delta$ temp | 3 | -104.88 | 215.78 | 0.74 | 0.09 |
| discharge + min discharge | 2 | -106.02 | 216.05 | 1.01 | 0.08 |
| discharge + min discharge + temp | 3 | -105.12 | 216.26 | 1.22 | 0.07 |
| min discharge + $\Delta$ temp | 2 | -106.16 | 216.33 | 1.29 | 0.07 |
| min discharge + temp | 2 | -106.16 | 216.33 | 1.29 | 0.07 |
| discharge + min discharge + $\Delta$ temp | 3 | -105.16 | 216.33 | 1.30 | 0.07 |
| discharge + $\Delta$ discharge + min discharge | 3 | -105.51 | 217.04 | 2.00 | 0.05 |
| $\Delta$ discharge + min discharge + temp + $\Delta$ temp | 4 | -104.51 | 217.06 | 2.02 | 0.05 |
| discharge + $\Delta$ discharge + min discharge + temp | 4 | -104.80 | 217.63 | 2.59 | 0.04 |
| discharge + $\Delta$ discharge + min discharge + $\Delta$ temp | 4 | -104.88 | 217.79 | 2.75 | 0.03 |
| discharge + min discharge + temp + $\Delta$ temp | 4 | -105.10 | 218.23 | 3.19 | 0.03 |
| min discharge + temp + $\Delta$ temp | 3 | -106.16 | 218.33 | 3.29 | 0.03 |
| discharge + $\Delta$ discharge + min discharge + temp + $\Delta$ temp | 5 | -104.45 | 218.94 | 3.90 | 0.02 |
| min discharge | 1 | -108.64 | 219.29 | 4.25 | 0.02 |

Model averaging of the 90% confidence sets found that in ‘early’ out-migrants, minimum discharge was significantly related to sturgeon beginning downriver migrations, but the confidence intervals for all other variables spanned 0 and, therefore, they were considered non-significant effects (Table 4). Green sturgeon likelihood of departure was positively related to minimum discharge values within a 5-d time interval (Hazard ratio = 1.48). Among green sturgeon that adopted the ‘early’ out-migration strategy, fish were more likely to depart at higher minimum discharge values—the mean minimum daily flows for the 5-d intervals when fish began their downriver migration was 218 m^3^ s^-1^ (range 118 to 396 m^3^ s^-1^).

**Table 4.** Model-averaged Cox proportional hazard parameter estimates for green sturgeon classified into either ‘early’ or ‘late’ out-migration groups in the Sacramento River.

| Covariate | Coefficient ($\beta$) &<br>Confidence Interval | S.E. $\beta$ | Hazard Ratio<br>(HR: $e^{\beta}$) | HR Confidence<br>Interval |
| --- | --- | --- | --- | --- |
| **(a) ‘Early’ out-migration** |  |  |  |  |
| Min discharge | 0.39 (0.14 – 0.65) | 0.13 | 1.48 | 1.15 – 1.92 |
| $\Delta$ discharge | -0.74 (-2.36 – 0.44) | 0.75 | 0.48 | 0.11 – 1.55 |
| $\Delta$ temp | -0.10 (-1.75 – 1.06) | 0.42 | 0.90 | 0.40 – 1.55 |
| Mean temp | -0.24 (-10.85 – 9.11) | 2.70 | 0.79 | 0.58 – 1.56 |
| Mean discharge | 0.02 (-1.09 – 1.19) | 0.39 | 1.02 | 0.48 – 2.18 |
| **(b) ‘Late’ out-migration** |  |  |  |  |
| Min discharge | 0.04 (0.01 – 0.07) | 0.02 | 1.03 | 1.00 – 1.07 |
| $\Delta$ discharge | -0.02 (-0.06 – 0.03) | 0.02 | 0.98 | 0.94 – 1.03 |
| $\Delta$ temp | -0.001 (-0.85 – 0.85) | 0.43 | 0.99 | 0.43 – 2.33 |
| Mean temp | 0.96 (-6.55 – 8.46) | 3.83 | 2.61 | 0.001 – 4.72 |
| Mean discharge | -0.006 (-0.04 – 0.02) | 0.01 | 0.99 | 0.96 – 1.02 |

Most green sturgeon migration events (68%) were classified into the ‘late’ out-migration group. Based on CPH model averaging, the timing of out-migrations in this group was related to the minimum discharge levels (HR = 1.03), but based on confidence intervals was not related to other variables (Table 4). Changes in discharge during the period of ‘late’ out-migrations included seasonal influxes of water that increased magnitude and variability in discharge, and also included a decline in the overall minimum discharge rate, with a 33% decline in minimum discharge from 187 ± 24 m^3^ s^-1^ at the beginning of the ‘late’ period to 126 ± 25 m^3^ s^-1^ over a period of 150 days. Considering both the 22-day profiles (Fig. 3) and CPH results, green sturgeon adopting a ‘late’ out-migration timing were likely to depart after initial increases in seasonal discharge rates, but also when minimum discharge levels were higher.

### Repeatability of migration timing

There was a positive correlation between the journey days that individuals began upriver migrations for their first and second observed migrations (Pearson’s correlation, t = 2.07, df = 19, P = 0.05; Fig. 4a), but there was not a corresponding correlation between journey days that downriver migrations began (Spearman’s correlation, rho = 0.29, P = 0.20; Fig. 4b). There was no correlation for the journey days that fish began migrating upriver and the day they began to move back downriver, i.e., individuals that migrated upriver earlier in the spring run did not necessarily return to the Pacific Ocean earlier (Spearman’s correlation, rho = -0.09, P = 0.56; Fig. 4c).

**Figure 4.**
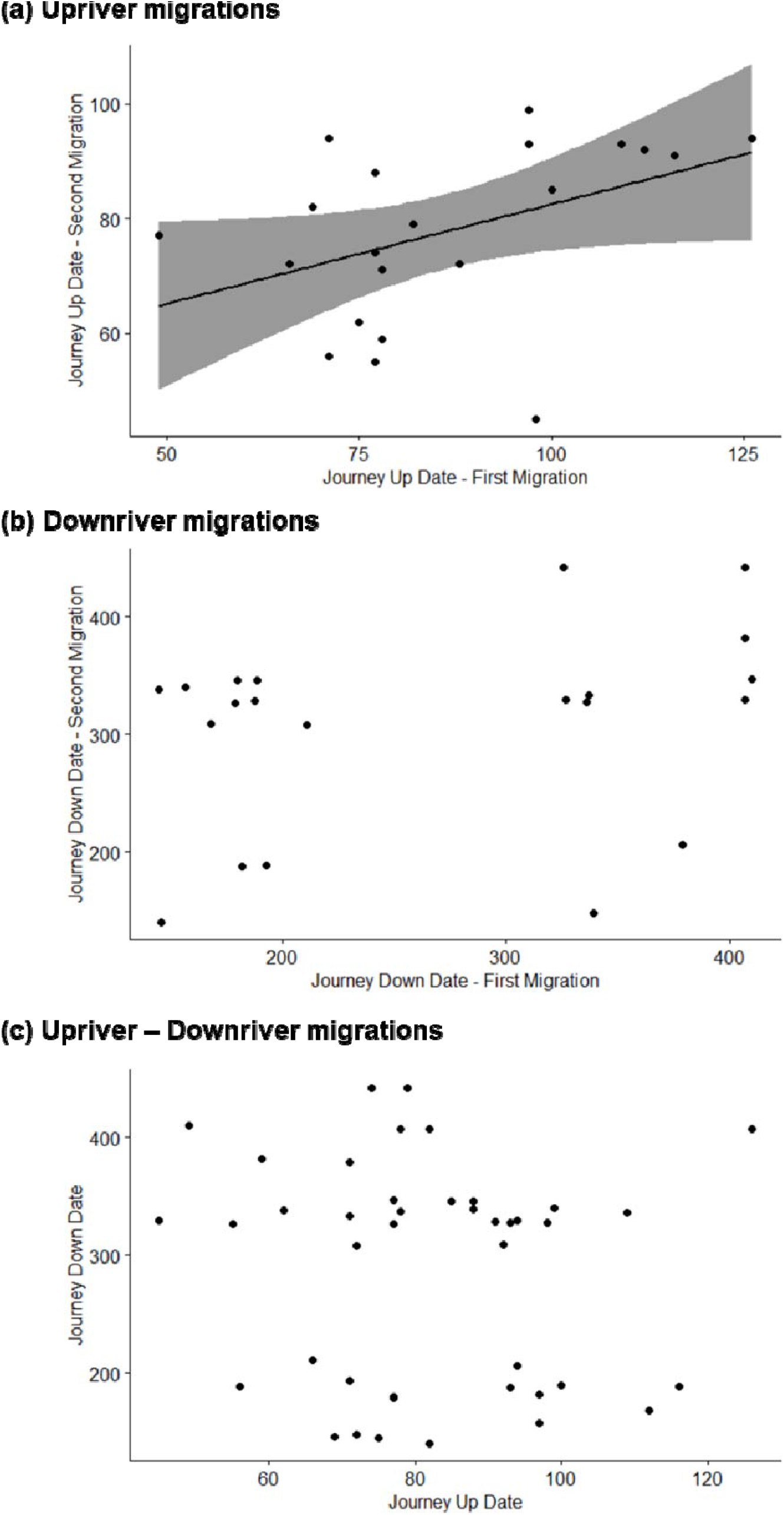
Comparison of migration timings (journey days) for individual green sturgeon that were tracked making more than one complete migration during the study observation period. Correlations of (a) upriver migration days, (b) downriver migration days, and (c) upriver day and the corresponding downriver date for a given migration are shown. Line of best fit is shown for correlations determined to be statistically significant. Only individuals with complete detection records for a given year from the time of entry to the Sacramento River were considered, i.e., fish tagged mid-migration in the river system were not included in these correlations because upriver journey days could not be determined.

Comparing swim down classifications between the first and second swim down events, 10 sturgeon adopted an ‘early-early’ strategy as compared to 13 with an ‘early-late’ strategy, representing 56% of fish changing from ‘early’ to ‘late’ timing between their first and second migrations (Fig. 5). In comparison, 33 fish adopted a ‘late-late’ strategy, with 8 fish (20%) changing strategy between migration periods (i.e., a ‘late-early’ strategy). Across all years of the observation period, we documented a total of 21 fish switching out-migration groups between their first and second migration intervals (13 fish early to late and 8 fish late to early) compared to a mean of 28 switches expected under random assignments over 100,000 simulations. The proportion of fish that switched strategy in these randomizations was found to be marginally significantly greater than the number observed empirically (P = 0.05). We interpreted this as a tendency for conservation of out-migration timing group between subsequent migrations, and further investigated the conditions under which a fish might switch strategy.

**Figure 5.**
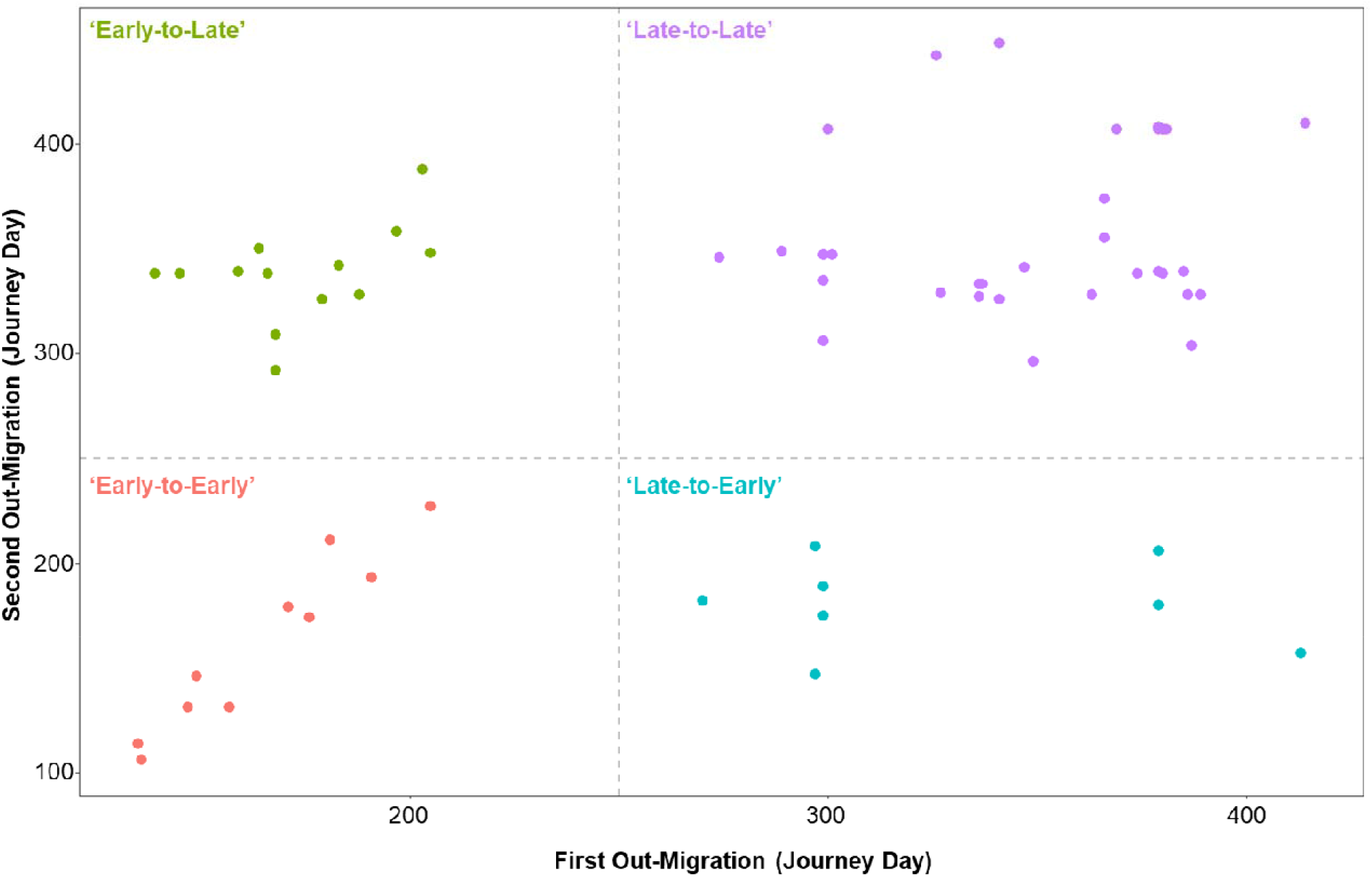
Timing of swim down for fish observed making two downriver migrations during the 2006-2018 observation period. Dashed lines represent the journey day 250 cut-off used to classify early and late swim down groups. For fish tagged mid-migration in the Sacramento River system the swim down journey date is given assuming they began to migrate upriver during the calendar year of capture.

Among the ‘early-early’ green sturgeon, there was a strong linear relationship between first and second departure journey days (Second migration day = 1.685 × First migration day – 116.47; R^2^ = 0.92), but on average fish departed 4 days earlier on their second observed migration. This estimated relationship was used to get matched dates for switching fish (i.e., ‘early-to-late’ and ‘late-to-early’, see Methods for details) and comparisons of mean flow during the ‘early’ departure period in years fish departed ‘early’ vs. holding to the ‘late’ departure period. Among the fish that switched migration timing (n = 21), river discharge was greater for the migration event when fish departed in the ‘early’ group (mean ± 1 S.E.: 283 ± 24 m^3^ s^-1^, median: 250 m^3^ s^-1^) as compared to the discharge levels matched to the ‘early’ period on years when fish departed ‘late’ (mean ± 1 S.E.: 187 ± 10 m^3^ s^-1^, median: 189 m^3^ s^-1^; Wilcoxon rank sums, Z = 205, P = 0.001). Therefore, among fish observed switching migration strategy (n = 23), sturgeon were more likely to depart during the ‘early’ period when the Sacramento River discharge rates were higher.

## Discussion

Biodiversity is currently declining across terrestrial and aquatic environments, but the rate of diversity loss among freshwater organisms is outpacing that of other major systems (40). For many species, conservation efforts have been fundamentally constrained by large gaps in the understanding of complete life-cycles and how species interact with habitats impacted by humans. Anadromous fishes that periodically enter freshwater rivers for spawning are emblematic of such challenges because (1) they spend much of their lifetime in ocean environments where they are rarely observed, (2) they migrate through a diverse array of habitats upon entering freshwater rivers, and (3) iteroparous species are infrequently observed across the multiple migration journeys they may make in a single lifetime. Green sturgeon in California are a prime example of how conservation management of endangered or threatened species must proceed even when we lack detailed information on the life-history and behavioral ecology of the species. In this study, we provide an example of how long-term biotelemetry studies can be collated to reveal complexity in behavioral life-histories and describe ecological characteristics of species directly relevant to conservation actions. Understanding the diversity and function of intraspecific life-history variability has implications for the management of species and ecosystems.

### Timing of migration events

The unimodal upriver migration of green sturgeon into the Sacramento River during the spring months, peaking during the month of March, resembled observations of the nDPS green sturgeon in Oregon (23). The discrete period of spring migrations for green sturgeon was consistent with selection pressures related to offspring development and survival (e.g., Wright and Trippel 2009, Tillotson and Quinn 2018). Reproductive success of fish migrating long distances from their home ranges to spawning grounds (such as sDPS green sturgeon migrating to California from areas near Vancouver Island; 43) may be under selection to ensure that arrival coincides with that of potential mates and optimal conditions for offspring development (44).

All the tagged green sturgeon returned to the Pacific Ocean with none of the tagged fish exhibiting permanent residency in the Sacramento River system. Miller et al. (22) reported that green sturgeon were detected in the Sacramento River system during all months of the year, raising the potential that the sDPS green sturgeon population includes partial migration strategies, i.e., some individuals exhibit permanent river residency. River-residents have been described for other sturgeon species, including lake sturgeon (45–47) and shortnose sturgeon (*Acipenser brevirostrum*; 48). Furthermore, alternate migration strategies are a common aspect of salmonid biology in California, presumably as a bet-hedging adaptation for the notoriously variable Mediterranean climate (49–51). Even though some of the green sturgeon monitored in this study spent over a year in the Sacramento River, they all eventually returned to the Pacific Ocean and on average returned to the river on a 4-year cycle (Supplemental Materials Fig. S1). The extended post-spawning holding time for some fish, i.e., the ‘late’ downriver group identified here, contributed to overlap among upriver and downriver migration groups across years, which explains the observation of green sturgeon present in the Sacramento River during all months (22). Given that some fish were present in the Sacramento River for over one year, there were likely foraging and habitat requirements specific to this residence period which are worthy of further investigation.

### Environmental cues of downriver migration

Heublein et al. (21) reported conflicting annual patterns relating discharge to downriver migrations across study years; these results may be due in part to the use of daily mean discharge and limited total sample size during initial tracking efforts. Indeed, mean discharge was considered during CPH model selection, but ranked low in relative importance scoring of variables (≤ 0.46 in both ‘early’ and ‘late’ groups). River flow characteristics have previously been described as likely drivers of green sturgeon migration (21). Our analysis across multiple years and repeated spawning events provided further support for river discharge as a primary factor influencing out-migration behaviors, particularly minimum discharge rates as measured across multiple days.

In addition to identification of the role minimum discharge for both ‘early’ and ‘late’ migrants in this study, seasonal patterns of flow variation in the system add context inferences of sturgeon migrations. ‘Early’ out-migrants were largely exiting during early summer months (June) and based on discharge profiles of the Sacramento River (Supplemental Materials Fig. S2), departed prior to annual lows in river discharge. While there has been interest in fish stranding occurring due to modifications of waterways, e.g., hydroelectric-related alterations to water discharge rates, there are natural sources of discharge variation that can also pose risks to fishes (reviewed by 52). It is possible that evolutionary responses to these natural cycles in river discharge resulted in cues observed here for the ‘early’ fish to depart the system at a perceived threshold or wait until levels increased during the winter months (‘late’ group).

In contrast to discharge patterns in the early summer months, the ‘late’ downriver group initiated migration during a period of seasonal discharge increases, and based on the 22-day profiles began out-migration after an influx of water. The pattern of an initial influx triggering downriver migration has also been reported for the nDPS green sturgeon (23) and was predicted for sDPS green sturgeon (53). Fish have been tracked moving downriver in response to changing discharge levels associated with stochastic events, including striped bass observed egressing the Hudson River before large-scale storm systems resulted in flow surges (54). Recent studies on high discharge rates and green sturgeon in the Sacramento River system have focused on factors impacting fish migrating upriver (55), but seasonal fluctuations in discharge may also influence evolution of differential out-migration patterns within this population.

Green sturgeon migration patterns have been predicted to be related to water temperature (23). In this study, we found links between discharge rates but not temperature for either out-migration group identified. Temperature is likely to have impacts on many aspects of green sturgeon life history, but it does not appear to be a primary factor in the migration patterns for sDPS green sturgeon (see Supplemental materials for additional discussion about temperature). Temperature may have a greater impact in rivers further north, e.g., nDPS population of green sturgeon in Oregon and Washington states, but the sDPS green sturgeon did not show patterns of water temperature predicting out-migration timing.

### Repeatability of migration timing across bouts

Multiple downriver migration groups with timing similar to those in this study have been reported for nDPS green sturgeon in the Klamath and Trinity rivers (23). In the Sacramento River system, the ‘late’ downriver group represented most observed migration events (68%), but the ‘early’ downriver departures still represented a significant portion of the downriver movements (32%). ‘Late’ departing fish were more likely to depart a second time in the same out-migration timing group than the ‘early’ departing fish (80% repeatability compared to 44%), nonetheless still nearly half of the ‘early’ group departed a second time in the ‘early’ group.

It has been speculated that rapid spring out-migration could be the result of tagging and handling effects due to previous observations of white sturgeon abandoning spawning runs following tagging (56). However, a previous study reported that 71% of green sturgeon tagged following stranding events in the Sacramento River continued moving upriver after tagging and release (55). In this study, we observed 62 repeat migration events separated by an average of 4 years and still observed the two distinct out-migration periods; therefore, we suggest ‘early’ and ‘late’ groups described here are unlikely to be related to tagging or handling effects on green sturgeon.

Repeatability of downriver migration times across spawning bouts was consistent with differential migration in sDPS green sturgeon. Many individuals did switch strategies, but overall, we observed a tendency of maintaining out-migration timing between subsequent events, and we argue that these tactics could be in part condition-dependent life history tactics rather than fixed for a lifetime. Life history variation is particularly of interest in fishes because it is widespread across taxa and falls into both fixed and conditional strategies (57). For example, Coho salmon (*Oncorhynchus kisutch*) males exhibit both morphologically distinct jack vs. hooknose life histories, fixed strategies maintained for the entire lifetime, and fight vs. sneak behaviors, conditional strategies that can change across reproductive events (58). Here, the presence of distinct downriver migration groups in green sturgeon may represent aspects of reproduction-related life history variation but given the ability of fish to switch groups across spawning bouts, this variation is likely a form of conditional life history strategy that can vary between reproductive bouts. Indeed, among the 23 fish observed switching between ‘early’ and ‘late’ strategies, we observed patterns in river discharge that support further consideration of river flow as the likely mechanism behind conditional switching of out-migration strategies. Our results comparing flow at early departures with flows at time-matched ‘early’ dates in years when fish out-migrated late, suggested the possibility that fish have a preferred ‘early’ departure times which they abandon in favor of a ‘late’ departure when flows are too low during their preferred ‘early’ time. Though we emphasize data only support this as a hypothesis and further work is needed, the possibility has intriguing implications for conservation and river flow management if months-delayed departures cause additional consequences ranging from energetic costs, particularly for post-spawn females, to risk of mortality.

### Anthropogenic impacts on life history variation

Anthropogenic impacts on communities may not impact all stages or forms of life history equally. For green sturgeon, the Red Bluffs Diversion Dam (RBDD) had a long period of potential impacts on sDPS migration from the mid-20^th^ century until its full deactivation in 2013 (53, 59). For much of its operational history, water gates of RBDD would have been closed during early summer months when the ‘early’ group would be migrating downriver. If these gates increased risk—either through injuries or mortality—to this specific group of fish it could account for some of the disparity observed between the overall number of fish adopting ‘early’ and ‘late’ migrations, especially for a long-lived intermittently spawning species such as green sturgeon.

Increased flows have been associated with increased spawning efficiency of diverse native fishes in California (60, 61), and much focus has been placed on the management of flow rates to increase spawning success and larval sturgeon survival (reviewed by 20). However, in this study we also found evidence that adult migration timing may be related to flow characteristics of the Sacramento River and that when fish experienced lower discharge rates during the late spring months (‘early’ group here) they may be more likely to hold over for several months in the river. Prior to its closure, the RBDD controlled water flow to 283 – 425 m^3^ s^-1^ (May - Sep) (62), which were consistent with the flow rates observed to be correlated with ‘early’ downriver migration here, but the dam itself presented a barrier to fish movements.

Among the fish that switched between ‘early’ and ‘late’ out-migration timing, we observed lower flow rates in the years fish switched to a ‘late’ strategy (mean = 187 m^3^ s^-1^), suggesting that facilitating adult green sturgeon migrations during the spring and early summer months will require more than deactivation of RBDD. Furthermore, if persistent drought conditions continue to impact river discharge, it is plausible that increasing numbers of green sturgeon will adopt a ‘late’ out-migration strategy. This extended river-holding behavior raises further questions about holding locations in the river, resource needs during the extended river holding period, physiological impacts of extended periods in freshwater, and if holding in the river increases exposure to other threats, e.g., susceptibility to poaching and overall capture risk.

## Conclusions

Our synthesis drew on telemetry data gathered for over a decade and provided further details on migration timing, identified potential environmental cues to downriver migration, and described within-population life history variability for the threatened population of green sturgeon in the Sacramento River. Long-term biotelemetry data therefore holds great potential for understanding the life-histories of species like sturgeon that conduct large-scale migrations (63, 64). Intraspecific variation in life histories can take many forms within wild populations, but ecological diversity and behavioral plasticity within populations may also provide buffering capacity to stochastic events. Individual-based tracking techniques, including acoustic telemetry, are providing avenues to describe and explore mechanisms of life history variation. These data are in turn valuable to conservation science aimed at protecting rare and declining species, like sturgeon. We concluded that the two downriver out-migration groups were robust across time and represented differential migration patterns based on the timing of movements, and we encourage their inclusion in conservation planning for sDPS green sturgeon. Furthermore, given the duration of activity in the Sacramento River and the potential for continued drought-related conditions causing stress on this populations, we recommend further examination of movement and habitat use within the upper reaches of the Sacramento River because adult green sturgeon may face stressors and risks in the river environment that could impact individual fitness and survival beyond the spawning season alone.

## List of Abbreviations

AIC: Akaike’s information criterion
CITES: Convention on International Trade in Endangered Species of Wild Fauna and Flora
CPH: Cox proportion hazard regression model
ESA: Endangered Species Act
HR: hazard ratio (part of CPH above)
nDPS: northern distinct population segment of green sturgeon
RBDD: Red Bluffs Diversion Dam
sDPS: southern distinct population segment of green sturgeon
VIF: variance inflation factor

## Declarations

### Availability of data and materials

All data regarding green sturgeon movements used in this study was compiled from the University of California Davis biotelemetry database (BARD - http://cftc.metro.ucdavis.edu/biotelemetry-autonomous-real-time-database/fishtrack).

### Authors’ contributions

SFC and LWS compiled all detection data and performed primary analysis and writing on the manuscript. DRO, DCR, JAW, and ALR provided feedback on all steps of analysis and manuscript drafts. GPS, JTK, and MT provided contextual support regarding green sturgeon in the Sacramento River system and comments on various drafts of this work. All authors have read and approved the final manuscript.

## Acknowledgements

The authors would like to recognize the contributions of many people in the UC Davis Biotelemetry Laboratory across multiple individual research projects that made this extensive green sturgeon dataset a possibility. We especially recognize A.P. Klimley for his leadership and efforts in organizing the initial ‘core 69 kHz array’, and CDFW for funding it. The number of people involved with these projects across multiple agencies and institutions are too numerous to list here but we express our sincerest gratitude for their dedication and hard work over many years.

