## Supplemental Table 1 for "Intraspecific variation in migration timing of green sturgeon in the Sacramento River system"

*Temperature and downriver movements*

Temperature was not related to the probability of green sturgeon beginning downriver migrations in either the ‘early’ or ‘late’ groups identified here. In the Klamath and Trinity rivers, part of the nDPS green sturgeon population, out-migration was observed when water temperatures were 10 – 12°C (Benson et al. 2007), comparable to the 11°C temperature for the ‘late’ group in this study, but lower than the 15°C for the ‘early’ group migrating in the early summer months. Information on the thermal tolerances and preferences of adult green sturgeon is sparse relative to early life history stages (reviewed by Rodgers et al. 2020), but tracking studies have detected green sturgeon presence across a wide range of temperatures in the San Francisco Estuary (14.5-20.8°C; Kelly et al. 2007) and along the coast of Oregon (9.5-16.0°C; Huff et al. 2012), well within the temperature ranges for the migration groups reported here.

Thermal tolerances for green sturgeon eggs, larvae, and juveniles have been thoroughly studied (reviewed by Rodgers et al. 2020), and the thermal optimum and tolerance ranges for these early life history stages is narrower than that of adults: the optimal range is 15-19°C for juveniles (Mayfield and Cech 2004). As such, adult timing to reach the spawning grounds over a relatively short window may be related to the more restrictive temperature requirements of eggs and early life history phases during development. In comparison, the out-migration back downriver occurred over a wider range of temperatures and was not related to temperature in the CPH models. The downriver return phase of migration would be related only to the thermal tolerances and preferences of adults, which may explain the wider range of temperatures over which the out-migration occurred.

Temperature is recognized as an important factor environmental variable for fishes in the Sacramento River, including and previous work has suggested a target to a goal of maintaining temperatures below 18°C to support early life stage development (Israel and Klimley 2008), but for adult green sturgeon in the Sacramento River, downriver migrations are likely also driven by flow characteristic, including discharge rates.

*Synthesis studies and biotelemetry databases*

When academic research groups, private companies, and government agencies co-ordinate the types of technology used, share infrastructure and maintenance requirements, and share collected data, the scope of projects can increase (Reubens et al. 2019). As such, tracking networks that share resources, workloads, and data have been formed on regional (e.g., Great Lakes Acoustic Telemetry Observation System, GLATOS, and Florida Atlantic Coast Telemetry Network, FACT), national (e.g., Integrated Marine Observing System, IMOS, in Australia), and international scales (e.g., Ocean Tracking Network, OTN, and European Tracking Network, ETN) (Krueger et al. 2018, Reubens et al. 2019, Young et al. 2020). As these networks grow and age, they have also begun to generate longitudinal detection databases providing new opportunities for synthesis-based studies and expanding the spatial scale of observations beyond those possible with many individual studies (Boucek and Morley 2019). We believe that this green sturgeon study is an example of how long-term, collaborative biotelemetry efforts are enhancing our understanding of basic ecology for wild populations and providing information applicable to specific conservation and management efforts. We encourage future efforts that leverage telemetry data across discrete projects to examine longitudinal patterns in movements and spatial ecology.

**Table S1.** Model fit summary for combinations of five covariations related to water discharge and temperature predicted to be related to the migration timing of green sturgeon assigned to the ‘early’ out-migration period in the Sacramento River system. Predictor variables for each model are shown along with the number of parameters in each model (*K*), log likelihood (*LL*), Akaike’s Information Criterion (AIC), difference in AIC score compared to the top model (*ΔAIC*), and model weight (*w_i_*).

| **Model variables** | ***K*** | ***LL*** | ***AICc*** | ***ΔAIC*** | ***w_i_*** |
| --- | --- | --- | --- | --- | --- |
| Δ discharge + min discharge | 2 | -29.52 | 63.06 | 0 | 0.19 |
| Δ discharge + min discharge + Δ temp | 3 | -28.93 | 63.92 | 0.86 | 0.12 |
| Δ discharge + min discharge + temp | 3 | -29.00 | 64.07 | 1.01 | 0.11 |
| discharge + Δ discharge + min discharge | 3 | -29.14 | 64.35 | 1.28 | 0.10 |
| discharge + min discharge | 2 | -30.39 | 64.80 | 1.74 | 0.08 |
| discharge + Δ discharge + min discharge + Δ temp | 4 | -28.62 | 65.34 | 2.28 | 0.06 |
| discharge + Δ discharge + min discharge + temp | 4 | -28.65 | 65.40 | 2.34 | 0.06 |
| discharge + min discharge + Δ temp | 3 | -29.70 | 65.46 | 2.40 | 0.06 |
| discharge + min discharge + temp | 3 | -29.82 | 65.69 | 2.63 | 0.05 |
| Δ discharge + min discharge + temp + Δ temp | 4 | -28.87 | 65.84 | 2.78 | 0.05 |
| discharge + min discharge + temp + Δ temp | 4 | -29.54 | 67.17 | 4.10 | 0.02 |
| min discharge | 1 | -32.65 | 67.31 | 4.24 | 0.02 |
| discharge + Δ discharge + min discharge + temp + Δ temp | 5 | -28.62 | 67.38 | 4.32 | 0.02 |
| min discharge + Δ temp | 2 | -32.04 | 68.11 | 5.05 | 0.02 |
| min discharge + temp | 2 | -32.13 | 68.29 | 5.23 | 0.01 |
| min discharge + temp + Δ temp | 3 | -31.94 | 69.93 | 6.87 | 0.01 |
| discharge + Δ discharge | 2 | -33.48 | 71.00 | 7.94 | 0.004 |
| discharge + Δ discharge + temp | 3 | -32.56 | 71.17 | 8.11 | 0.003 |
| discharge + Δ discharge + Δ temp | 3 | -32.61 | 71.27 | 8.21 | 0.003 |
| discharge + Δ discharge + temp + Δ temp | 4 | -32.55 | 73.19 | 10.13 | 0.001 |
| Δ temp | 1 | -37.15 | 76.31 | 13.25 | 0.0003 |
| discharge + Δ temp | 2 | -36.18 | 76.39 | 13.33 | 0.0002 |
| discharge + temp | 2 | -36.29 | 76.62 | 13.56 | 0.0002 |
| temp | 1 | -37.31 | 76.63 | 13.57 | 0.0002 |
| discharge | 1 | -37.79 | 77.59 | 14.53 | 0.0001 |
| Δ discharge + Δ temp | 2 | -36.80 | 77.62 | 14.56 | 0.0001 |
| Δ discharge + temp | 2 | -36.94 | 77.91 | 14.85 | 0.0001 |
| temp + Δ temp | 2 | -37.10 | 78.22 | 15.16 | 0.0001 |
| discharge + temp + Δ temp | 3 | -36.14 | 78.33 | 15.27 | 0.0001 |
| Δ discharge + temp + Δ temp | 3 | -36.73 | 79.53 | 16.47 | 0.0001 |
| Δ discharge | 1 | -39.05 | 80.11 | 17.05 | 0.0000 |

**Table S2.** Model fit summary for combinations of five covariations related to water discharge and temperature predicted to be related to the migration timing of green sturgeon assigned to the ‘late’ out-migration period in the Sacramento River system. Predictor variables for each model are shown along with the number of parameters in each model (*K*), log likelihood (*LL*), Akaike’s Information Criterion (AIC), difference in AIC score compared to the top model (*ΔAIC*), and model weight (*w_i_*).

| **Model variables** | ***K*** | ***LL*** | ***AICc*** | ***ΔAIC*** | ***w_i_*** |
| --- | --- | --- | --- | --- | --- |
| Δ discharge + min discharge | 2 | -105.51 | 215.04 | 0 | 0.13 |
| Δ discharge + min discharge + temp | 3 | -104.80 | 215.62 | 0.58 | 0.10 |
| Δ discharge + min discharge + Δ temp | 3 | -104.88 | 215.78 | 0.74 | 0.09 |
| discharge + min discharge | 2 | -106.02 | 216.05 | 1.01 | 0.08 |
| discharge + min discharge + temp | 3 | -105.12 | 216.26 | 1.22 | 0.07 |
| min discharge + Δ temp | 2 | -106.16 | 216.33 | 1.29 | 0.07 |
| min discharge + temp | 2 | -106.16 | 216.33 | 1.29 | 0.07 |
| discharge + min discharge + Δ temp | 3 | -105.16 | 216.33 | 1.30 | 0.07 |
| discharge + Δ discharge + min discharge | 3 | -105.51 | 217.04 | 2.00 | 0.05 |
| Δ discharge + min discharge + temp + Δ temp | 4 | -104.51 | 217.06 | 2.02 | 0.05 |
| discharge + Δ discharge + min discharge + temp | 4 | -104.80 | 217.63 | 2.59 | 0.04 |
| discharge + Δ discharge + min discharge + Δ temp | 4 | -104.88 | 217.79 | 2.75 | 0.03 |
| discharge + min discharge + temp + Δ temp | 4 | -105.10 | 218.23 | 3.19 | 0.03 |
| min discharge + temp + Δ temp | 3 | -106.16 | 218.33 | 3.29 | 0.03 |
| discharge + Δ discharge + min discharge + temp + Δ temp | 5 | -104.45 | 218.94 | 3.90 | 0.02 |
| min discharge | 1 | -108.64 | 219.29 | 4.25 | 0.02 |
| discharge + Δ discharge + temp | 3 | -106.83 | 219.67 | 4.63 | 0.01 |
| discharge + Δ discharge + Δ temp | 3 | -107.01 | 220.04 | 5.00 | 0.01 |
| discharge + temp | 2 | -108.17 | 220.34 | 5.30 | 0.01 |
| discharge + Δ temp | 2 | -108.23 | 220.46 | 5.43 | 0.01 |
| discharge + Δ discharge + temp + Δ temp | 4 | -106.28 | 220.59 | 5.55 | 0.01 |
| Δ discharge + temp | 2 | -109.11 | 222.23 | 7.19 | 0.004 |
| discharge + temp + Δ temp | 3 | -108.15 | 222.32 | 7.29 | 0.003 |
| Δ discharge + Δ temp | 2 | -109.19 | 222.38 | 7.34 | 0.003 |
| temp | 1 | -110.72 | 223.44 | 8.40 | 0.002 |
| discharge + Δ discharge | 2 | -109.84 | 223.69 | 8.65 | 0.002 |
| Δ temp | 1 | -111.10 | 224.21 | 9.17 | 0.001 |
| Δ discharge + temp + Δ temp | 3 | -109.10 | 224.21 | 9.17 | 0.001 |
| temp + Δ temp | 2 | -110.33 | 224.66 | 9.62 | 0.001 |
| discharge | 1 | -113.37 | 228.74 | 13.70 | 0.0001 |
| Δ discharge | 1 | -115.48 | 232.97 | 17.93 | 0.00002 |

**Table S3.** Relative importance scores for each covariate modelled in relation to (a) ‘early’ and (b) ‘late’ out-migration timing of green sturgeon from the Sacramento River system. Models are ordered to present relative importance scores in the same order as models are presented in Tables S2 and S3 (see above).

| **Group** | **Discharge** | **Min discharge** | **Δ discharge** | **Temperature** | **Δ Temperature** |
| --- | --- | --- | --- | --- | --- |
| ***‘Early’ out-migration*** | | | | | |
|  | 0.022 | 0.022 | 0.022 | 0.022 | 0.022 |
|  | 0.000 | 0.190 | 0.000 | 0.115 | 0.000 |
|  | 0.004 | 0.123 | 0.000 | 0.047 | 0.123 |
|  | 0.003 | 0.115 | 0.190 | 0.000 | 0.047 |
|  | 0.100 | 0.100 | 0.123 | 0.000 | 0.000 |
|  | 0.061 | 0.080 | 0.115 | 0.059 | 0.000 |
|  | 0.059 | 0.061 | 0.047 | 0.003 | 0.003 |
|  | 0.003 | 0.059 | 0.000 | 0.001 | 0.061 |
|  | 0.001 | 0.057 | 0.000 | 0.051 | 0.001 |
|  | 0.000 | 0.051 | 0.004 | 0.024 | 0.000 |
|  | 0.080 | 0.047 | 0.003 | 0.000 | 0.057 |
|  | 0.057 | 0.024 | 0.100 | 0.000 | 0.024 |
|  | 0.051 | 0.023 | 0.061 | 0.014 | 0.000 |
|  | 0.024 | 0.015 | 0.059 | 0.006 | 0.015 |
|  | 0.000 | 0.014 | 0.003 | 0.000 | 0.006 |
|  | 0.000 | 0.006 | 0.001 | 0.000 | 0.000 |
| **Sum** | **0.466** | **0.987** | **0.729** | **0.344** | **0.361** |
| ***‘Late’ out-migration*** | | | | | |
|  | 0.019 | 0.019 | 0.019 | 0.019 | 0.019 |
|  | 0.000 | 0.132 | 0.000 | 0.099 | 0.003 |
|  | 0.002 | 0.099 | 0.003 | 0.048 | 0.091 |
|  | 0.011 | 0.091 | 0.132 | 0.004 | 0.048 |
|  | 0.048 | 0.079 | 0.091 | 0.001 | 0.001 |
|  | 0.033 | 0.071 | 0.099 | 0.036 | 0.001 |
|  | 0.036 | 0.069 | 0.048 | 0.013 | 0.011 |
|  | 0.013 | 0.069 | 0.004 | 0.008 | 0.033 |
|  | 0.008 | 0.069 | 0.001 | 0.071 | 0.008 |
|  | 0.009 | 0.048 | 0.002 | 0.027 | 0.009 |
|  | 0.079 | 0.048 | 0.011 | 0.003 | 0.069 |
|  | 0.069 | 0.036 | 0.048 | 0.009 | 0.027 |
|  | 0.071 | 0.033 | 0.033 | 0.069 | 0.003 |
|  | 0.027 | 0.027 | 0.036 | 0.025 | 0.069 |
|  | 0.003 | 0.025 | 0.013 | 0.002 | 0.025 |
|  | 0.009 | 0.016 | 0.008 | 0.001 | 0.001 |
| **Sum** | **0.438** | **0.932** | **0.548** | **0.436** | **0.420** |

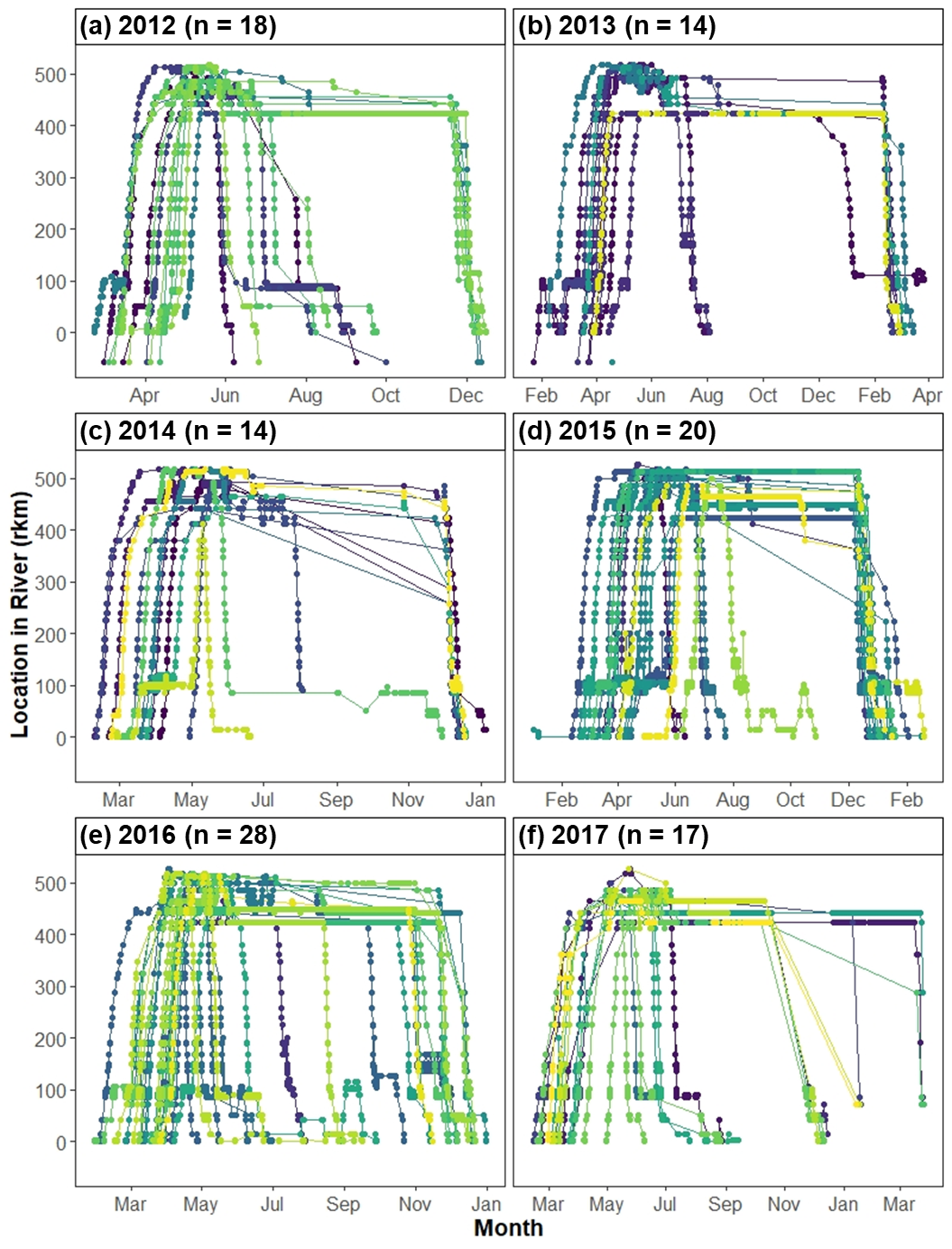

**Figure S1.** Migration profile plots displaying the River km (rkm) associated with each detection of all green sturgeon commencing a presumed spawning run for the years 2012 through 2017. Green sturgeon detections were examined across the years 2006 – 2018, but only years with > 10 individuals detected migrating both up and downriver are shown here. Colors represent the migration profile tracks through the river for individual green sturgeon.

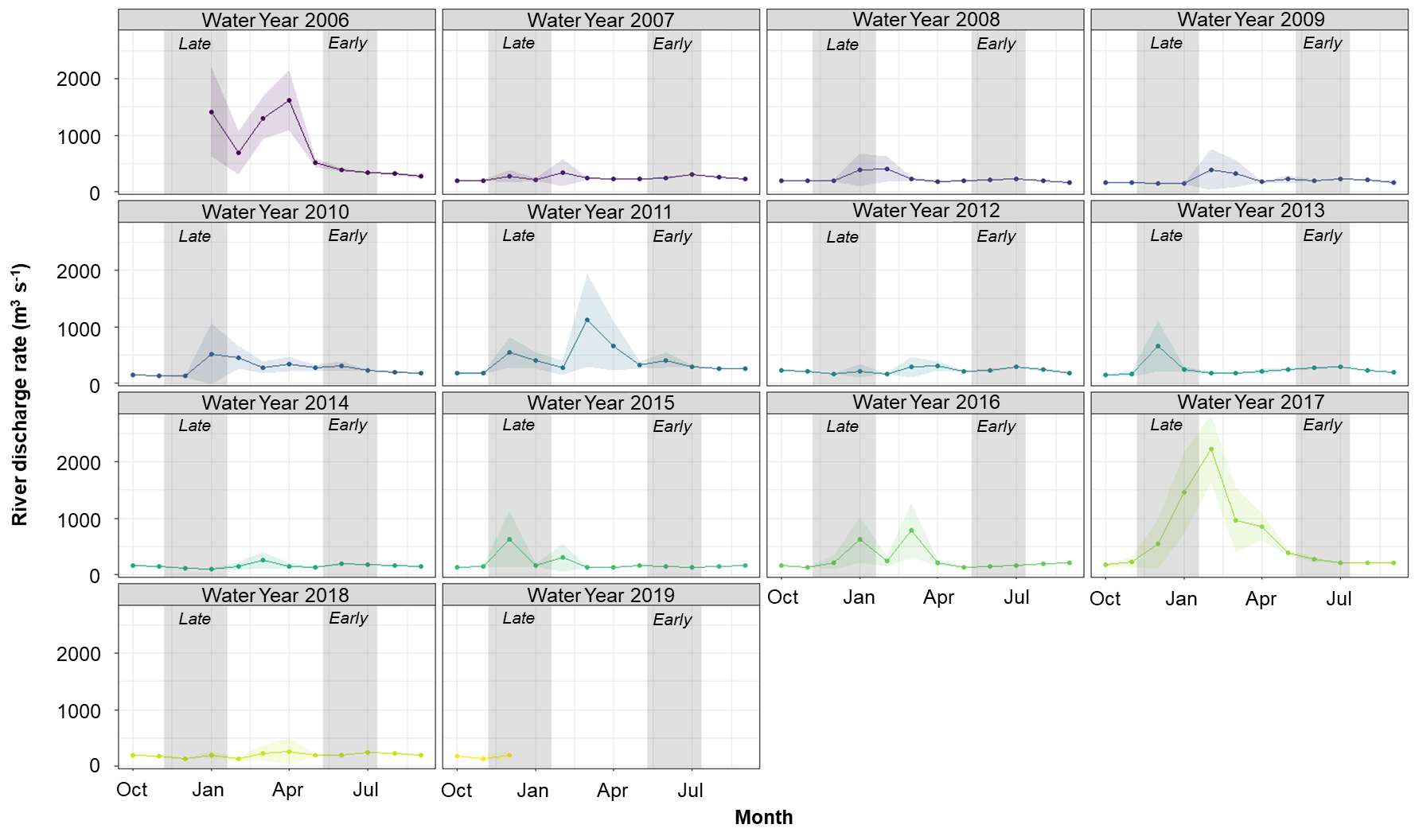

**Figure S2.** Sacramento river discharge rates (m^3^ s^-1^) as measured in the upper portion of the river (ORD station) based on California water year calendar dates (beginning in October of each year). Monthly average discharge is shown with ribbons reflecting 1 S.E. and boxes represent the period of ‘early’ and ‘late’ swim down events across all years.

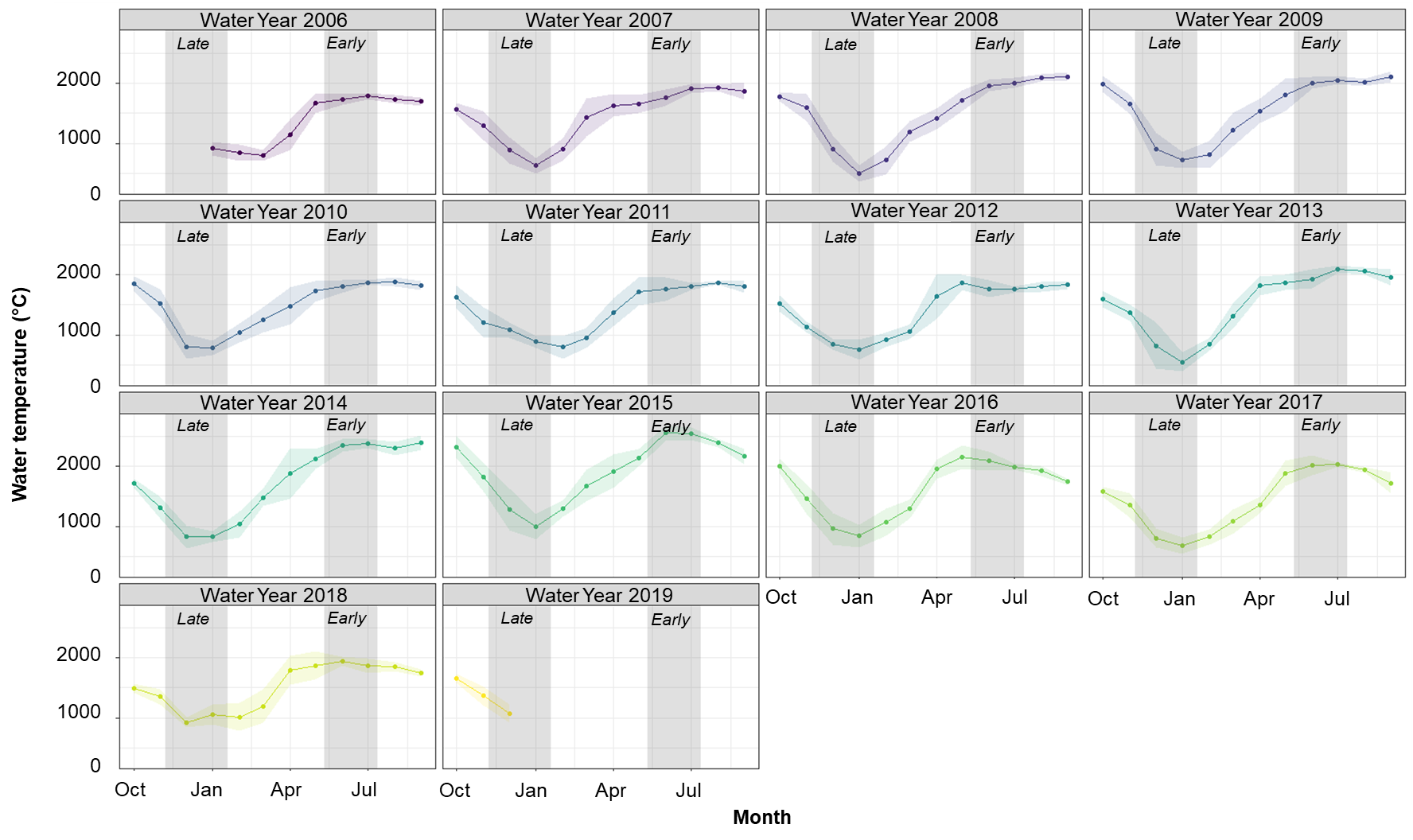

**Figure S3.** Sacramento river water temperature (°C) as measured in the upper portion of the river (RDB station) classified into California water years (i.e., beginning 1 October of each year). Monthly average temperature is shown with ribbons reflecting 1 S.E. and boxes represent the period of ‘early’ and ‘late’ swim down events across all years.
